# GSEA-InContext Explorer: An interactive visualization tool for putting gene set enrichment analysis results into biological context

**DOI:** 10.1101/659847

**Authors:** Rani K. Powers, Anthony Sun, James C. Costello

**Affiliations:** Computational Bioscience Program, University of Colorado Anschutz Medical Campus, Aurora, CO, USA; Department of Pharmacology, University of Colorado Anschutz Medical Campus, Aurora, CO, USA; Colorado School of the Mines, Golden, CO, USA; University of Colorado Cancer Center, University of Colorado Anschutz Medical Campus, Aurora, CO, USA

**Author notes:** **Corresponding author:** James C. Costello, University of Colorado Anschutz Medical Campus, 12801 E. 17th Ave. Mail Stop 8303, Aurora, CO 80045.

## Abstract

**Summary:** GSEA-InContext Explorer is a Shiny app that allows users to perform two methods of gene set enrichment analysis (GSEA). The first, GSEAPreranked, applies the GSEA algorithm in which statistical significance is estimated from a null distribution of enrichment scores generated for randomly permuted gene sets. The second, GSEA-InContext, incorporates a user-defined set of background experiments to define the null distribution and calculate statistical significance. GSEA-InContext Explorer allows the user to build custom background sets from a compendium of over 5,700 curated experiments, run both GSEAPreranked and GSEA-InContext on their own uploaded experiment, and explore the results using an interactive interface. This tool will allow researchers to visualize gene sets that are commonly enriched across experiments and identify gene sets that are uniquely significant in their experiment, thus complementing current methods for interpreting gene set enrichment results.

**Availability and implementation:** The code for GSEA-InContext Explorer is available at: https://github.com/CostelloLab/GSEA-InContext_Explorer and the interactive tool is at: http://gsea-incontext_explorer.ngrok.io

## 1) Introduction

Gene Set Enrichment Analysis (GSEA) (Mootha *et al.*, 2003; Subramanian *et al.*, 2005) is one of the most widely used methods in bioinformatics for interpreting coordinate, pathway-level changes in -omics experiments. Functionally similar genes are grouped into gene sets, then tested for enrichment in the positive or negative direction against an individual genome-wide experiment. By testing gene sets rather than individual genes, GSEA provides increased explanatory power, reduced complexity, and can offer insight into the underlying biology of dysregulated pathways and processes in the experiment being analyzed (Khatri *et al.*, 2012). For experiments with less than seven samples per condition, the recommended statistical model for calculating significance is GSEAPreranked (GSEA User Guide, 2018), which estimates *p*-values using an empirical null distribution of enrichment scores generated on permuted gene sets, wherein genes are randomly selected from the input experiment.

Previously, we reported a meta-analysis of GSEAPreranked results for 442 transcriptome experiments in which cancer cell lines were treated with various small molecules (Powers *et al.*, 2018). We found that many gene sets were consistently up or down regulated across experiments. For example, the TNFA_SIGNALING_VIA_NFKB gene set from the Hallmarks collection (Liberzon *et al.*, 2015) was significantly up-regulated in over 50% of the tested experiments (FDR < 0.05). Based on these observations, we developed a complementary method to GSEAPreranked called GSEA-InContext, which accounts for gene expression patterns based on a background set of experiments to identify statistically significantly enriched gene sets. In doing so, GSEA-InContext addresses a novel research question: which pathways are uniquely enriched in one experiment compared to many other independent experiments? Applying GSEA-InContext to different experiments, we demonstrated that our method was able to extract results that were biologically relevant and experiment-specific. We supplied GSEA-InContext as a Python package, along with experimental data and overall results. In order to make the GSEA-InContext algorithm more accessible to researchers, here we present a Shiny web application: GSEA-InContext Explorer (http://gsea-incontext_explorer.ngrok.io), which includes a user-friendly web interface to interactively build background sets from over 5,700 experiments, run GSEA-InContext and GSEAPreranked, and visualize how a user’s experiment of interest compares to the compendium of other experiments.

## 2) Materials and methods

### 2.1) Data

In total, 5,718 experiments were compiled from three separate sources. First, 442 experiments were compiled as reported in (Powers *et al.*, 2018). Human gene expression studies of cancer cell lines treated with a drug were downloaded from Gene Expression Omnibus. Experiments on the Affymetrix U133 plus 2.0 platform with at least 3 replicates and an appropriate vehicle control were processed using RMA (Bolstad *et al.*, 2003). Ranked gene lists were determined by the log-fold change of treatment compared to vehicle control. Second, 5,005 experiments were compiled from a data set of NCI-60 cell lines treated with 15 small molecule drugs (GSE116436) (Monks *et al.*, 2018); data were processed using the same workflow as in (Powers *et al.*, 2018). Third, 271 experiment were compiled from the Connectivity Map build 1 (GSE5258) (Lamb *et al.*, 2006), again following the workflow in (Powers *et al.*, 2018). All data, including sample-level annotations are available through the GSEA-InContext app.

### 2.2) GSEA-InContext Explorer workflow

The GSEA-InContext algorithm requires two inputs: 1) a “background set” of experiments that is used to inform the statistical testing procedure, and 2) a ranked list of genes from a user input gene expression experiment. We illustrate the analysis workflow using **Figure 1**.

**Figure 1.**
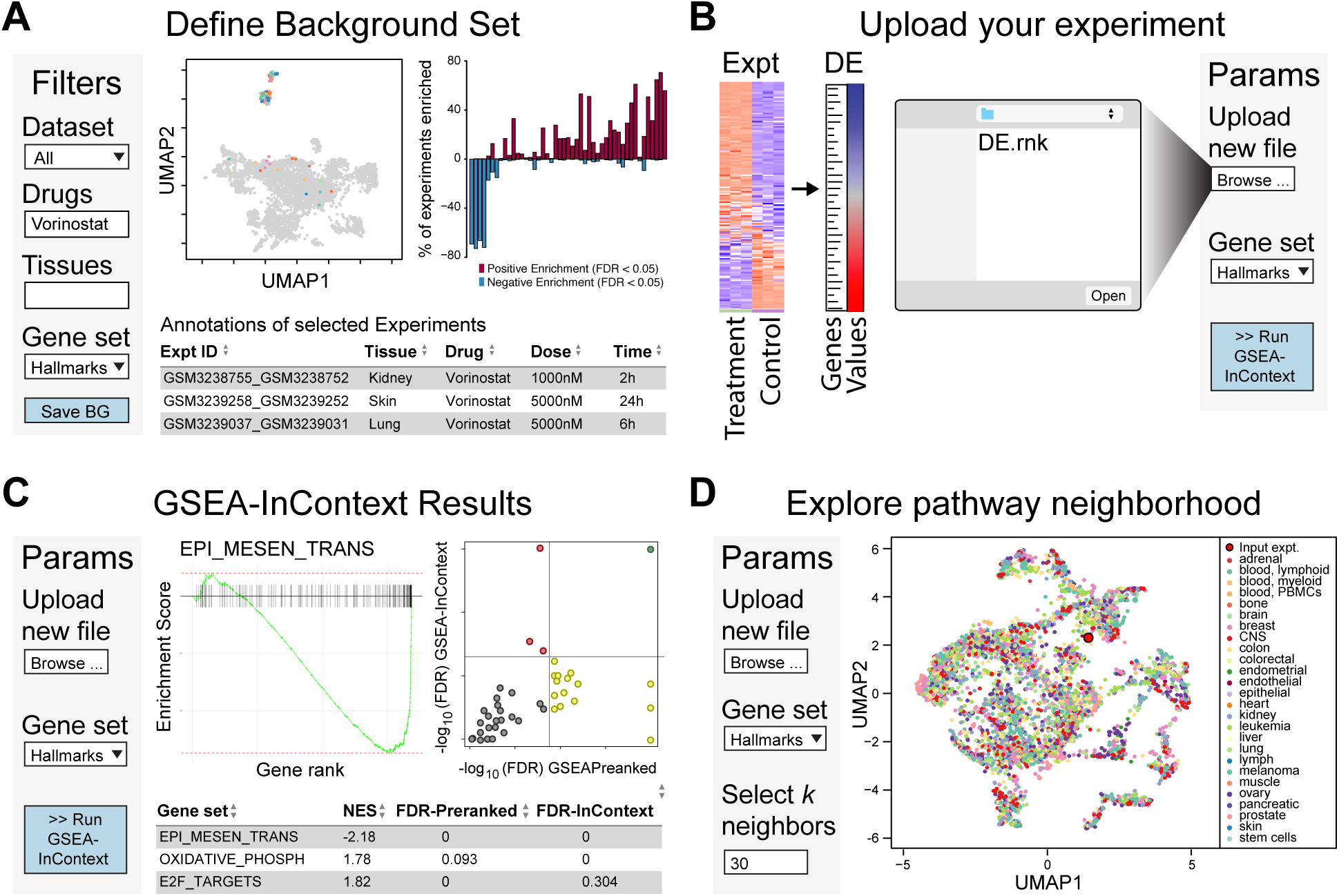
Workflow of GSEA-InContext Explorer. (**A**) The landing page of the app, where the user can define a background set of experiments based off of filters for dataset, drug, and tissue type. Selected experiments are displayed in the UMAP visualization of all 5,718 experiments. Vorinostat, an HDAC inhibitor, was queried and the associated experiments are colored according to cell type, with the remaining experiments greyed out. The associated barplot displays significant gene sets (FDR < 0.05) for the selected gene set collection (Hallmarks (Liberzon *et al.*, 2015)). After the user has defined a background set, clicking the “Save BG” button will save this set for use in GSEA-InContext. (**B**) A user can upload their differential expression file in .rnk format and then run GSEA-InContext. (**C**) Results of GSEAPreranked and GSEA-InContext are displayed, with a GSEA plot shown for any gene set selected from the table below. The FDR corrected *p*-values for both methods are displayed in a scatter plot. (**D**) The uploaded experiment will be projected onto the UMAP representation of all 5,718 experiments based on the normalized enrichment score of the selected gene set collection. The user can select the number of nearest neighbor experiments that will be listed in the table below.

#### 2.2.1) Background data set selection

**Figure 1A** illustrates the interface by which a user can query over 5,700 experiments to build an experiment-specific background set by setting filters for the desired platform, drug, tissue, etc. Two plots are displayed that dynamically update as the filter criteria are modified: a uniform manifold projection (UMAP) plot (McInnes *et al.*, 2018) computed on the ranked gene lists, and a barplot summarizing the results of GSEAPreranked analysis on the selected experiments for a gene set collection. After the appropriate background is established, the user can save the background set for use in the GSEA-InContext algorithm.

#### 2.2.2) Upload a gene expression experiment

**Figure 1B** shows that a user can upload their gene expression experiment in the .rnk format, which is a two column, tab-delimited text file where the first column is human gene symbols and the second is the differential expression values (often, log-fold change). The submitted file will be validated and then the GSEA-InContext algorithm can be executed using the previously defined background set and the selected gene set collection.

#### 2.2.3) GSEA-InContext analysis

**Figure 1C** illustrates the interface that displays the results of GSEAPreranked and GSEA-InContext, as reported in tabular form at the bottom of the app and visualized in a plot which compares the respective FDR corrected p-values. In this plot the gene sets are colored according to their significance as determined by the two methods: grey points are not significant in either method, green points are significant in both GSEAPreranked and GSEA-InContext, yellow points are only significant in GSEAPreranked, and red points are only significant in GSEA-InContext. The user can select any gene set from the table of gene set results to display the corresponding enrichment plot.

#### 2.2.4) Pathway context explorer

**Figure 1D** shows a novel functional exploration tool whereby the user’s input data can be projected onto a UMAP plot of the full set of over 5,700 experiments. The projection is based on the NES values from the gene set collection run in the GSEA-InContext analysis. The user can define the number of closest neighbors, which will populate the table underneath with the associated experiments selected from the interactive UMAP.

## 3) Discussion

The development of GSEA-InContext Explorer is motivated by the need for researchers to be able to compare their GSEA results to GSEA results obtained from other experiments. This allows researchers to address two key questions in relationship to their experiment: i) which gene sets are commonly enriched across a compendium of experiments, and ii) which gene sets are uniquely enriched in my experiment compared to many other independent experiments? Thus, the tool will allow researchers to complement and enrich the results generated by GSEA in order to ask deeper questions about their experiment.

In the future, we will add RNA-seq data, the ability for users to upload their own background data sets, and add user accounts for saving analysis results.

